# Identifying aligned tensor decompositions across heterogeneous neural datasets

**DOI:** 10.64898/2026.11.12.766278

**Authors:** Miklós Králik, Sasha Tinelli, James Kozloski, Rui Ponte Costa, Andrew Sharott, Robert L. Peach

## Abstract

Large-scale neural recordings increasingly span many sessions, brain regions and animals, but differences in neuronal sampling make it difficult to identify and interpret shared patterns. We introduce Neuralign, a tensor-based model that jointly aligns and decomposes recordings into components, each pairing a shared temporal pattern with its expression in individual trials and neurons, without requiring matched neurons, matched trials or behavioural labels. In simulations, pooling recordings improved recovery of shared components; in recurrent networks, Neuralign distinguished related from distinct task solutions. In the macaque motor cortex, label-free components recovered the ring-like organisation of reach directions. Across 93 brain-wide mouse recordings, Neuralign discovered a single component that isolated the response to inhibition of the anterior lateral motor cortex (ALM). A reference learned from correct, uninhibited trials revealed inhibition-related activity outside the choice direction on error trials, most strongly in the ALM and thalamus. Neuralign provides an interpretable framework for studying the shared structure of large neural datasets, linking shared patterns to individual trials and recorded populations.

## 1 Introduction

Advances in electrophysiology and optical imaging now allow the activity of thousands of neurons to be recorded simultaneously across multiple sessions, brain regions and animals, often in combination with behavioural tasks and experimental perturbations [1–4]. Analysing such recordings jointly could reveal activity patterns that recur across sessions and animals [5, 6], identify the differential expression of those patterns across neural populations, or even determine whether different animals or networks share a similar solution for a task. In practice, however, each recording samples a different set of neurons and usually contains a varying number of trials, making it challenging to both combine and decompose such data, ultimately limiting scientific insight.

Each shared activity pattern is characterised by its time course within the trial, its amplitude on individual trials and its distribution across neurons, and each of these can differ between recordings and sessions. A pattern associated with movement preparation, for example, may have the same time course in every session, whilst its amplitude varies from trial to trial with behaviour and perturbation, and its distribution across neurons and brain regions differs between animals. Interpreting multi-session data therefore requires a representation that establishes correspondence between recordings whilst retaining features of each shared pattern in every recording.

Population alignment methods establish correspondence between recordings by mapping their activity into a common low-dimensional space [7]. Canonical correlation analysis, Procrustes alignment, Hyperalignment and the shared-response model (SRM) learn a linear map for each recording, often using condition-averaged trajectories as matched observations [8–11]. These representations support comparisons of population geometry and the transfer of decoders between sessions. Non-linear and multi-session models extend this approach to single-trial dynamics [12], contrastive embeddings [13–15], the geometry of neural flow fields [16], and large pre-trained decoders [17, 18]. Related models combine session-specific neuronal weights with shared dynamics or temporal bases [15, 19]. However, a common space does not by itself provide an explicit description of when each recurring activity pattern occurs, how its strength varies across trials nor the contributions of individual neurons.

Tensor component analysis (TCA) provides this component structure for a single recording, decomposing a neuron × time × trial tensor into components that each combine a time course (temporal factor), an amplitude on every trial (trial factor) and a weight for every neuron (neuron factor). Despite these interpretable properties, to allow correspondence between components (up to an orthogonal transformation) TCA requires the same neurons to be tracked across sessions [20], which is difficult within an animal and not possible between animals. To overcome this, one can apply TCA after alignment, revealing variation across trials and sessions within a common population space [21], but the correspondence between recordings is then fixed before the components are estimated. Estimating the alignment and decomposition jointly would instead allow the components themselves to establish correspondence, whilst pooling evidence across recordings.

Here we introduce Neuralign, a tensor decomposition model that jointly aligns and decomposes trial-resolved neural recordings from different sessions and animals. Because recordings time-locked to the same task events share a time axis, each component has a single temporal factor shared across recordings, whereas its trial and neuron factors are estimated separately for each recording. The neuron factors are coupled across recordings through a shared component geometry, mapped onto the neurons of each recording by a session-specific projection and scaled by session gains. Inspired by PARAFAC2 and coupled tensor models [22–25], the coupling is imposed as a soft penalty, such that the neuron factors of each recording can retain session-specific variation. Neuralign thus pools evidence about shared activity patterns without matching individual neurons or trials and retains the factors needed to relate their expression to experimental variables. Neuralign is unsupervised by default, and labels such as reach target or rewarded and unrewarded trials can anchor the alignment without entering the decomposition. Besides the components, the fitted model provides a linear projection of recorded activity into the shared space, which can be restricted to neurons from a single brain region and applied to trials not used for fitting.

We applied Neuralign to four datasets, ordered by how much of the shared structure was known in advance, allowing us to first validate and then showcase its capability to generate novel insights into real data. First, in simulations with known generating factors, we tested whether fitting sessions jointly recovered shared components more accurately than TCA fitted to each session or alignment with SRM and Hyperalignment. In low-rank recurrent networks with known dynamical solutions, we tested whether differences between networks were retained in the session gains and trial factors after alignment. In the macaque motor cortex, we related shared components to reach direction, cortical area, animal identity and hand position. Finally, in 93 brain-wide mouse recordings that all included ALM but sampled other regions in different combinations, we used Neuralign without inhibition labels to identify the shared response to optogenetic inhibition of ALM, and used the aligned projection to locate the regions in which inhibition remained detectable on error trials. Together, these applications show how a single decomposition can address a wide and varying set of questions posed by multi-session data, through identifying shared patterns and relating them back to the neural populations, individuals and behaviour.

## 2 Results

### 2.1 Neuralign aligns and decomposes heterogeneous neural recordings

We designed Neuralign to estimate the latent structure that is shared between recordings but distributed differently across neurons and trials (Fig. 1). Each recording session is represented as a neuron × time × trial tensor *Z*^(*n*)^ where the time axis is shared across sessions, imposed by time-locking to a common task event, whilst the number of neurons and trials can vary across sessions (Fig. 1a,b). Neuralign decomposes all sessions jointly into *R* components, each pairing a temporal factor shared across sessions (*V*) with neuron and trial factors specific to each session (*A*^(*n*)^ and *U* ^(*n*)^; Fig. 1c). The temporal factors describe the expression of each component with time, the trial factors capture how strongly a component is expressed on each trial, and the neuron factors capture the contribution of each recorded neuron to the component (Fig. 1c–e). The shared temporal factors establish a common identity across recordings without requiring matched neurons or trials.

**Fig. 1.**
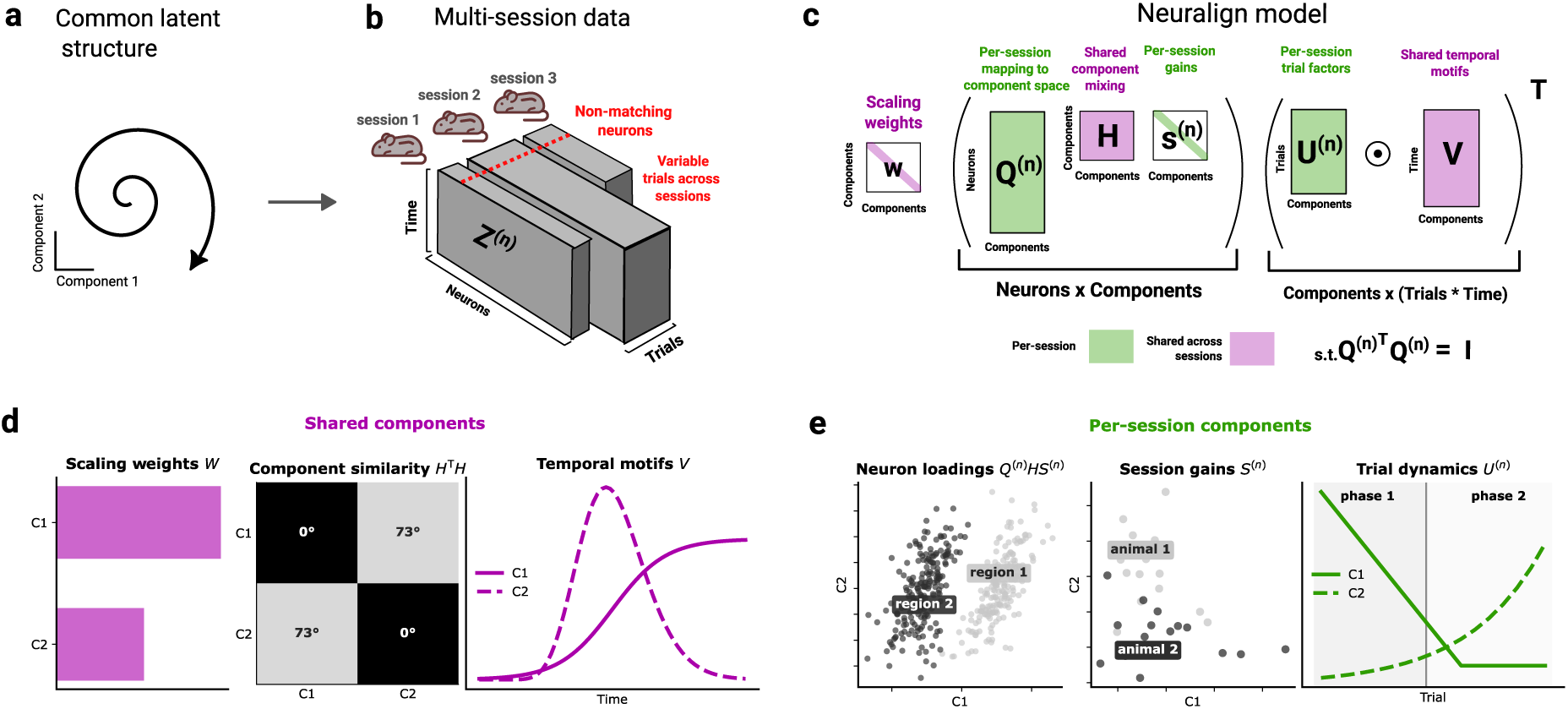
Neuralign links shared temporal activity patterns to neurons and trials across recordings. **a,b**, Neural recordings can share common latent dynamics. Recordings with different neurons and trial counts are organised as sets of neuron-by-time-by-trial tensors. **c**, Neuralign joint tensor decomposition: factors are coupled to a common component geometry while retaining session-specific variation. Magenta denotes shared quantities and green session-specific quantities. **d**, Shared components: shared component weights *w* set the global scale of the decomposition. The mixing matrix *H* defines the shared component geometry through pairwise component similarity. The temporal factors *V* specify shared neuronal activity patterns. **e**, Per-session components: neuron loadings *A*^(^*^n^*^)^ *≈ Q*^(^*^n^*^)^*H* diag(*S*^(^*^n^*^)^) represent individual neurons’ contributions to the component space, often clustering into structures such as regions. Session gains *S*^(^*^n^*^)^, which scale the strength of individual components, can reveal a meta-structure across the recordings, such as the experimental subjects. The trial factors *U* ^(^*^n^*^)^ reveal how component expression changes over the course of a recording session, such as a transition from one learning phase to the next.

In addition to the shared temporal factors, Neuralign couples the neuron factors across sessions through a shared component geometry (Fig. 1c). The neuron factors of each session are approximated as *A*^(*n*)^ ≈ *Q*^(*n*)^*H* diag(*S*^(*n*)^), where *Q*^(*n*)^ is a session-specific orthonormal projection, *H* is a shared mixing matrix and *S*^(*n*)^ contains the session gains. The mixing matrix defines the geometry of the components through their pairwise similarities (*H^T^ H*; Fig. 1d), the projection maps this geometry onto the neurons recorded in each session, and the session gains scale the expression of each component within that session (Fig. 1e). Differences in session gains can reveal structure across recordings, such as grouping by experimental subject. The coupling is imposed as a soft penalty, allowing the neuron factors to retain session-specific variation, and, together with the temporal factors, places all sessions in a common component space.

We fit Neuralign without condition labels by default (label-free Neuralign). In this setting, each session’s projection *Q*^(*n*)^ is estimated by orthogonal Procrustes, aligning the trial-averaged activity with the mean trajectory predicted by the model, whilst the reconstruction objective is computed on individual trials (Methods). When condition labels are available, such as reach target or rewarded/unrewarded trials, condition-averaged trajectories provide additional anchors for alignment (condition-aware Neuralign). Labels therefore guide the alignment of sessions without entering the reconstruction objective.

### 2.2 Pooling recordings improves recovery of shared factors

We first used simulated recordings, in which the generating factors are known, to test whether Neuralign recovers shared structure despite differences in neuronal sampling and session-specific activity, and whether this recovery improves as we increase the number of recording sessions. Each session contained two shared components, generated with the temporal factors and component geometry assumed by Neuralign, together with a weak session-specific (private) component and Poisson spike-count variability (Fig. 2a,b). The number of neurons and trials, the neuron loadings and session gains varied across sessions, with session gains drawn from two groups to mimic recordings from two animals. Neuralign was fitted at the known shared rank of two and compared with TCA fitted to each session separately, and with SRM and Hyperalignment fitted jointly across sessions (Methods). In an example cohort of 36 sessions, the temporal and trial factors recovered by Neuralign more closely matched the generating factors (Fig. 2c).

**Fig. 2.**
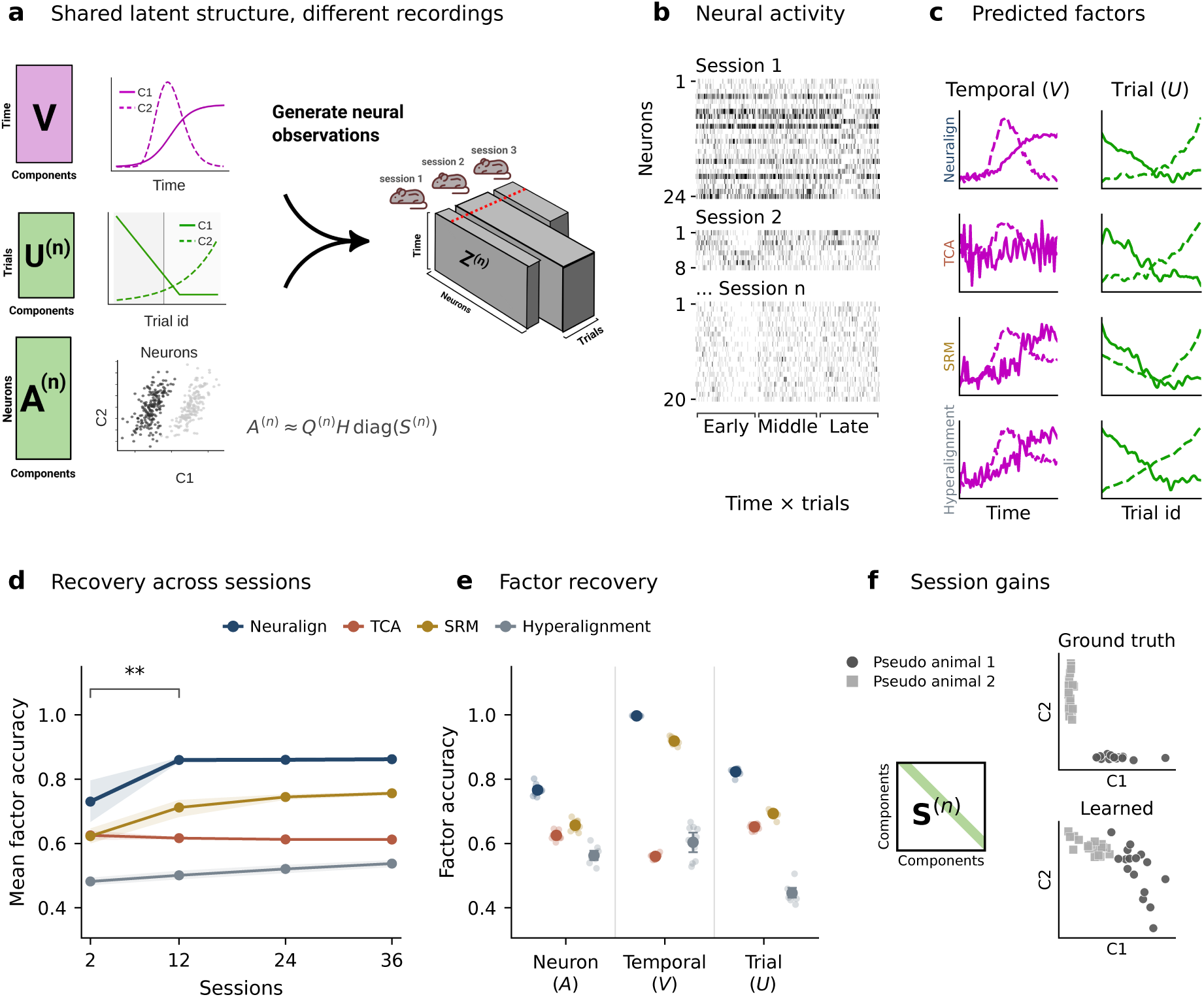
Pooling recordings improves recovery of shared factors. **a,b**, Two shared components and weak private activity generate recordings with different neurons and trial counts; greyscale heatmaps show example synthetic spiking activity. **c**, Recovered temporal and trial factors across decomposition methods, from a 36-session example, matched to ground truth and normalised for display. **d,e**, Factor-match scores as an increasing number of recording sessions are fit (**d**) and scores disaggregated by factor mode at 36 sessions (**e**). Lines or large points show means; shading and bars show bootstrap 95% confidence intervals across ten simulated datasets. Faint points show individual datasets. The bracket compares Neuralign at 12 versus 2 sessions (one-sided paired Wilcoxon test, *p* = 0.00684). **f**, Generating and learned session gains for two simulated gain groups. Components are matched and scaled for display.

Recovery was assessed with a factor-match score, which matches fitted factors to the generating factors and averages their absolute cosine similarity across the neuron, temporal and trial factors. A score of 1 indicates exact recovery up to order, sign and scale (Methods).

Across ten independently simulated datasets, the mean factor-match score of Neuralign increased from 0.730 with two sessions to 0.860 with 12 sessions, with little further improvement at 24 and 36 sessions (Fig. 2d). Neuralign exceeded the other methods at every cohort size, including TCA, whose per-session fits cannot benefit from additional recordings. Broken down by factors, the shared temporal factors were recovered almost exactly, and the remaining error lay within the neuron and trial factors, although their recovery still exceeded that of other methods (Fig. 2e).

The learned session gains also separated sessions from the two simulated groups (Fig. 2f), indicating that differences in component expression between recordings are retained rather than averaged away. Together, these simulation experiments suggest that pooling recordings improves recovery of shared factors whilst preserving differences in their expression across recordings.

### 2.3 Shared factors distinguish related from distinct low-rank RNN solutions

Having established recovery of shared factors, we next asked whether the shared representation also preserves differences between dynamical solutions, which alignment alone could obscure. Measuring similarity between systems that perform the same task, whether animals or trained networks, is a key use case of multi-session alignment models [6, 16]. Low-rank recurrent neural networks (RNNs) provide a controlled setting for this problem, because networks trained on the same task can converge to qualitatively distinct dynamical solutions whose structure is directly interpretable [26, 27].

Depending on the initialisation, rank-2 RNNs trained on a delayed match-to-sample task learn to encode the task in one of two topologically distinct attractor structures. We analysed two networks with the same structures (solutions I/A and I/B) and one with a distinct structure (solution II), each simulated across 20 input gains that scale the stimulus input (Fig. 3a–c). Using the experimental setup from MARBLE [16], we asked whether Neuralign could identify this graded similarity structure, i.e., two solutions that are topologically similar should be better aligned to each other than to a topologically distinct solution.

**Fig. 3.**
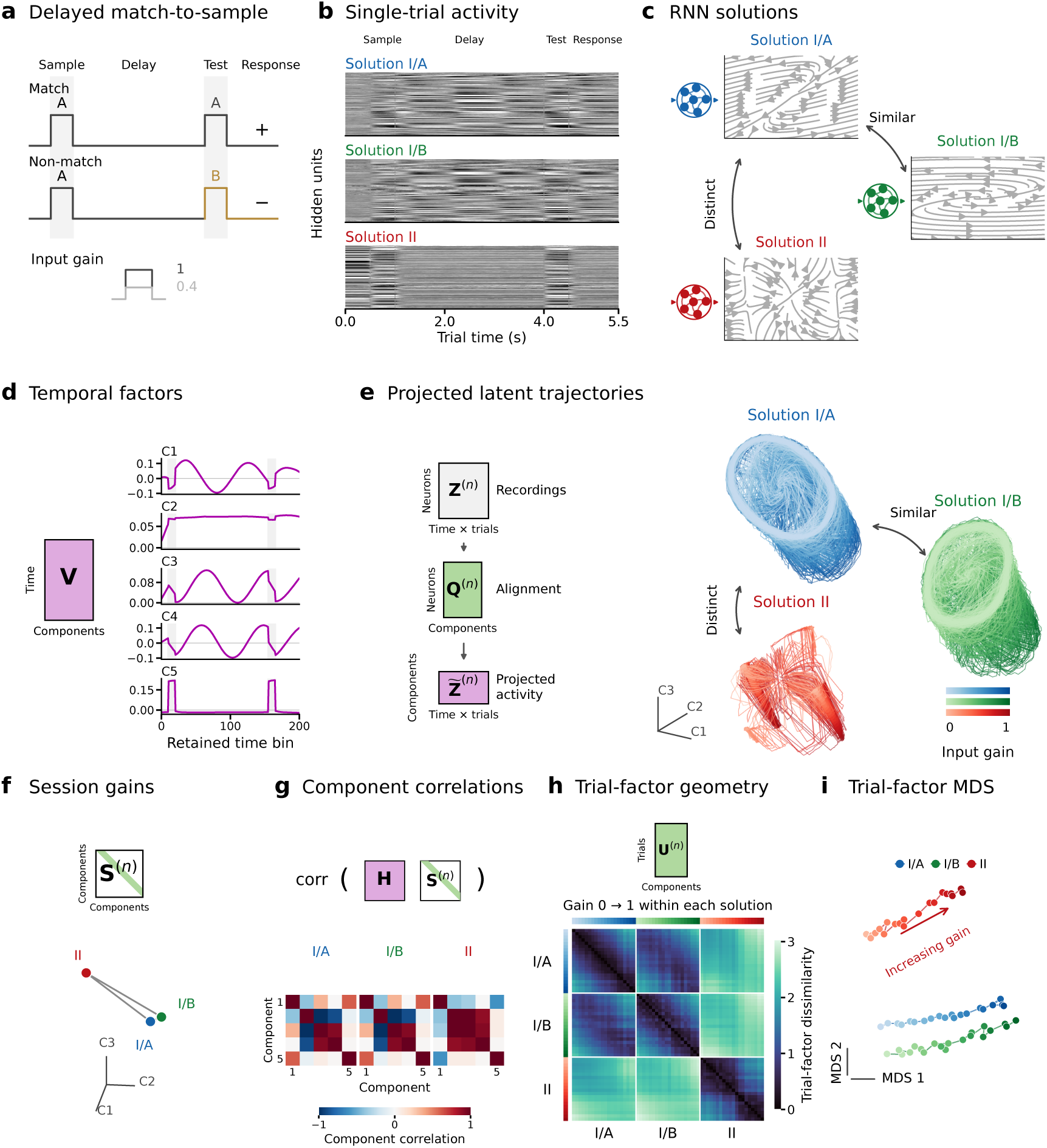
Shared factors distinguish related low-rank RNN solutions. **a**, Delayed match-to-sample task and input gain. **b**, One gain-1 trial per network, showing normalised activity of RNN hidden units. **c**, Intrinsic flow fields for two related solutions (I/A, I/B) and a distinct solution (II). **d**, All five shared temporal factors from the Neuralign decomposition; grey windows mark stimulus bins. **e**, Measured activity projected directly into the first three Neuralign coordinates, with a shared camera angle. Darker shades indicate higher input gain. **f**, Session gains from the first three Neuralign components, with equal axis scales and a connecting triangle. **g**, Full five-component Gram matrices *G*^(^*^n^*^)^ = diag(*S*^(^*^n^*^)^)*H^⊤^H* diag(*S*^(^*^n^*^)^), showing signed pairwise correlations between components for the different solutions. **h**, Wasserstein distances between all solution–gain distributions of the trial factors; lower values indicate greater similarity. Gain increases from 0 to 1 within each solution. **i**, MDS of the same distance matrix; lines join neighbouring gains.

We jointly fitted a label-free rank-5 Neuralign model to the hidden-unit activity of all three RNNs, treating each network as a separate session (see Supplementary Fig. S2a,b for rank selection). The shared temporal factors described recurring task-related patterns, including transient responses to the sample and test stimuli (Fig. 3d), and projecting each network’s activity into the common component space revealed similar trajectory geometries for I/A and I/B that differed from that of II (Fig. 3e). The session gains of I/A and I/B and the component correlations that combine these gains with the shared geometry *H* were likewise more similar to each other than to those of II (Fig. 3f,g). Trial-factor distributions, which describe how component expression varies across trials and input gains, were closer between I/A and I/B than between either network and II, as summarised by their pairwise Wasserstein distances and multidimensional scaling (MDS) embedding (Fig. 3h,i). Together, these results show that Neuralign can align non-linear systems while retaining interpretable differences between their dynamical solutions.

### 2.4 Shared motor components reveal reach geometry and improve kinematic decoding

We next applied Neuralign to recordings from the macaque motor cortex to examine how shared components relate to reach direction, brain region and movement kinematics. The dataset comprised 36 sessions from three macaques (C, M and T) performing an eight-target centre-out reaching task, with simultaneous recordings from primary motor cortex (M1) and dorsal premotor cortex (PMd) [28] (Fig. 4a,b). We fitted a label-free rank-6 Neuralign model to all sessions, without information about reach target, brain region or animal identity (rank selection in Supplementary Fig. S3e,f).

**Fig. 4.**
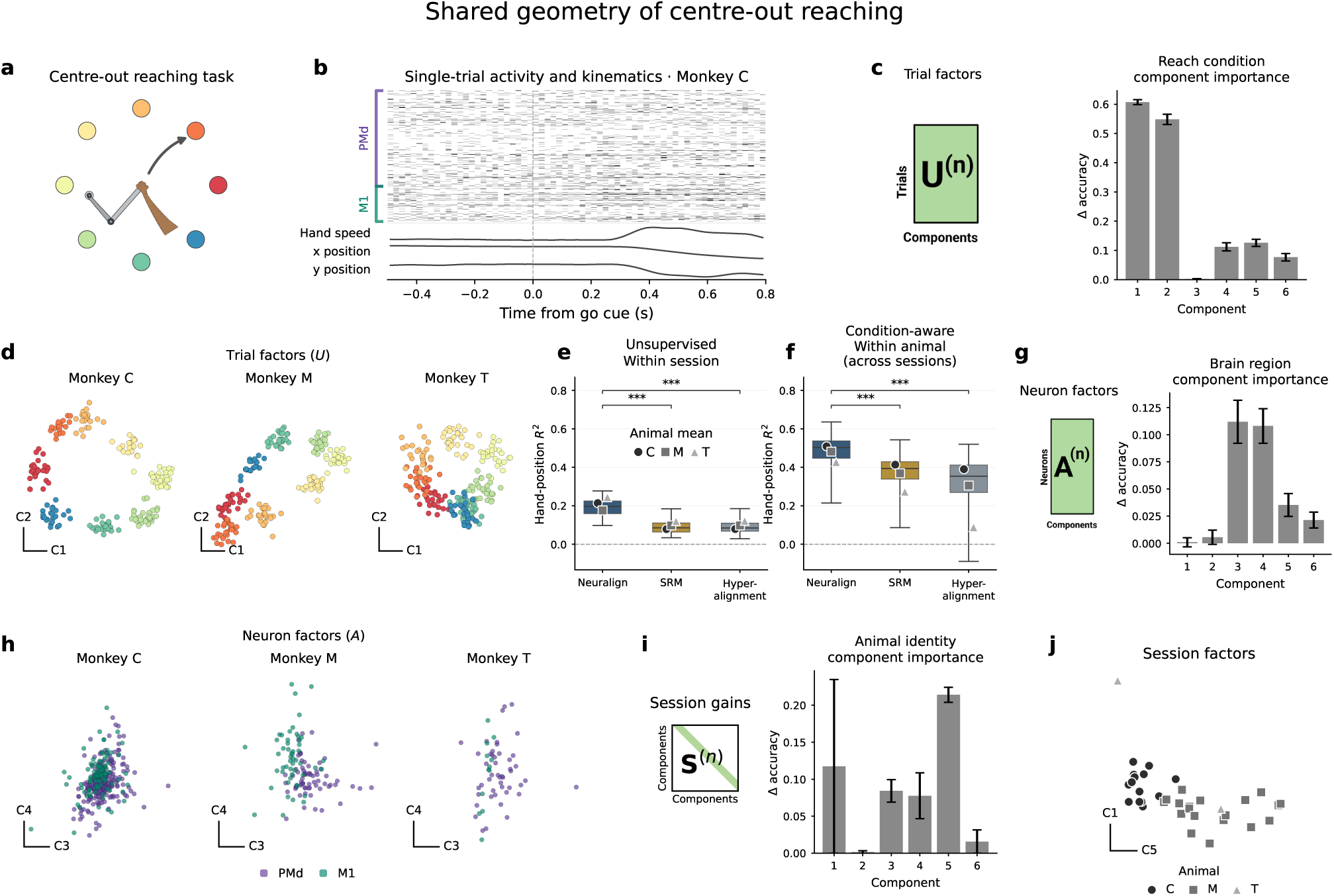
Motor components link task and neural geometries. **a,b**, Centre-out reaching task and a representative trial, showing regional spike counts, hand speed and position. Dashed line marks the go cue, which instructs the animal to move a two-link manipulandum, controlling a cursor on a screen, towards the instructed target. **c,d**, Trial-factor importance for reach target and the two most informative components in example sessions from each animal. **e,f**, Post-go hand-position decoding within sessions using label-free representations (**e**) and across sessions within an animal using condition-aware representations (**f**). In this figure only panel (**f**) uses a condition-aware Neuralign fit. Brackets mark one-sided paired Wilcoxon tests with BH correction (\*\*\**q <* 0.001). **g,h**, Neuron-factor importance for region and the two most informative components. **i,j**, Session-gain importance for animal identity and the two most informative gains. Importance bars show mean *±* s.e.m. across sessions in **c,g**, and decoder folds in **i**. All recordings contribute to representation fitting; held-out sessions refer to decoder training.

In an example session from each macaque, the trial factors of the two components most predictive of reach direction (C1 and C2; Fig. 4c) arranged trials in a ring that followed the circular layout of the targets (Fig. 4d). To test whether the representations also captured how the movement unfolded beyond its categorical reach target, we decoded hand position after the go cue from the representations of Neuralign, SRM and Hyperalignment (none of which were fitted using the kinematic data; Fig. 4e,f). Label-free Neuralign improved decoding within sessions (mean *R*^2^ = 0.195, compared with 0.092 for both SRM and Hyperalignment). When reach-target labels were provided to all methods to guide alignment, condition-aware Neuralign also improved decoding across sessions from the same animal (mean *R*^2^ = 0.489, compared with 0.381 (SRM) and 0.326 (Hyperalignment)).

Beyond decoding, Neuralign’s neuron factors showed a slight separation between M1 and PMd along components C3 and C4, the two components most informative about brain region (Fig. 4g,h). This indicated that these components were expressed in both regions but weighted differently between them, highlighting both mixed selectivity and regional specificity of the neural responses. In the label-free fit, the session gains also grouped recordings by animal (Fig. 4i,j), similar to our previous simulated examples (Fig. 2f). Thus, a ring-like reach geometry shared across animals coexisted with regional and animal differences in component expression, and the same representation improved the readout of movement kinematics.

### 2.5 A shared component captures inhibition-related activity across recordings

We next tested whether the Neuralign decomposition could recover the effect of optogenetic inhibition of the ALM across brain-wide electrophysiological recordings. In the Mesoscale Activity Map dataset [29], mice reported an auditory instruction with a delayed left or right lick. The ALM was simultaneously inhibited unilaterally or bilaterally during the late delay on a subset of trials, an intervention that can induce choice reversal (Fig. 5a,b).

**Fig. 5.**
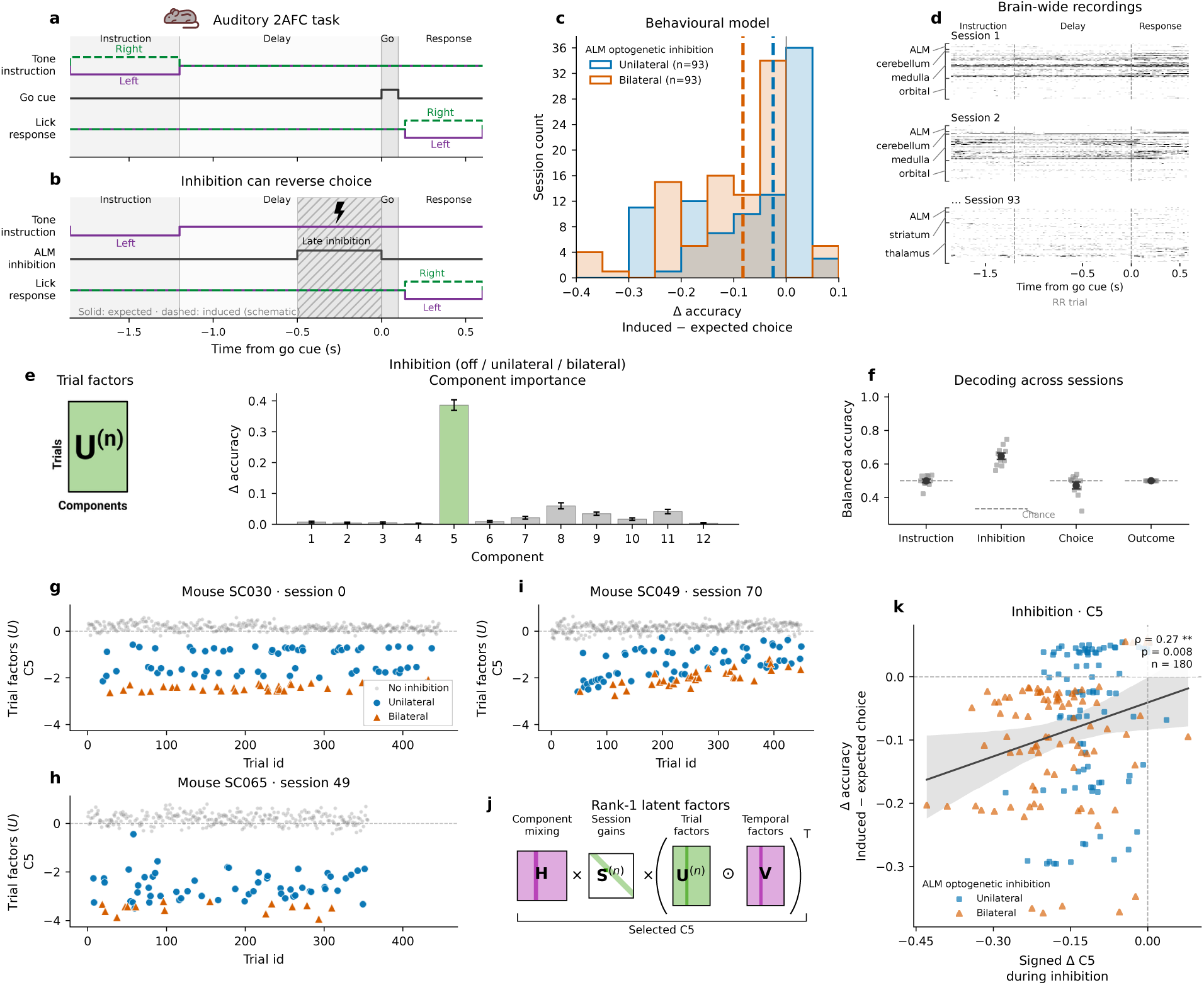
A shared component captures ALM inhibition-related activity across recordings. **a,b**, Delayed-lick auditory 2AFC (2-alternative forced choice) task and a schematic choice reversal during late-delay ALM inhibition. **c**, Histograms of behavioural linear mixed effects model-estimated accuracy changes across 93 sessions; negative values indicate impairment and dashed lines mark medians. **d**, Single-trial spike counts from three example sessions, grouped by region. **e**, Trial-factor schematic and inhibition importance across components (mean *±* s.e.m. across 87 eligible sessions); green highlights component 5. **f**, Decoding with one component selected separately (same protocol as in (**e**)) for each task feature within training folds, evaluated in ten held-out-session folds. Small points show folds; larger points and bars show mean *±* s.e.m.; dashed lines mark chance. Instruction (left/right tone); Inhibition (off/unilateral/bilateral); Choice (left/right); Outcome (hit/miss). **g–i**, Component-5 trial factors in three example mice: no inhibition (grey), unilateral (blue circles) and bilateral inhibition (orange triangles). **j,k**, Selected-component latent reconstruction and its signed change against behavioural accuracy change (*n* = 180 session–inhibition observations). The line and shading show a visualisation fit and session-bootstrap 95% confidence band. Reports Spearman correlation and a two-sided permutation test (\*\**p <* 0.01).

We first quantified how behavioural susceptibility to inhibition varied across recording sessions, using a mixed-effects model to estimate the change in correct-choice accuracy under inhibition relative to no inhibition, with negative values indicating poorer performance (Methods). Bilateral inhibition produced a larger median impairment than unilateral inhibition, and impairment varied widely across sessions (Fig. 5c), providing a per-session behavioural measure against which inhibition-related neural changes could be compared.

We then fitted a label-free rank-12 Neuralign model to all 93 sessions from 19 mice, allowing inhibition responses to be compared despite large differences in regional sampling (Fig. 5d; Supplementary Fig. S4c–e). Although inhibition labels were not used during fitting, the trial factors of a single component (C5) were by far the most informative about the three inhibition classes: no inhibition, unilateral inhibition and bilateral inhibition (Fig. 5e).

Using only the information from C5, a linear discriminant analysis (LDA) decoder classified inhibition in held-out sessions with a balanced accuracy of 0.647 ± 0.019 (mean ± s.e.m. over ten folds), compared with a chance level of 1*/*3 (Fig. 5f). By contrast, components selected in the same way for instruction, choice and outcome decoded these features near chance in held-out sessions. In example sessions from three mice, C5 trial factors were near zero on uninhibited trials and shifted away from zero on inhibited trials, more strongly under bilateral than unilateral inhibition (Fig. 5g–i). Under the same single-coordinate transfer protocol, rank-12 SRM and Hyperalignment did not isolate a comparably transferable inhibition (balanced accuracy 0.3542 ± 0.0034 and 0.3336 ± 0.0002, respectively; Supplementary Fig. S5f–i).

We assessed whether inhibition-related changes in C5 were associated with behavioural impairment by reconstructing the contribution of C5 in aligned latent coordinates and comparing inhibited with uninhibited trial means during the late delay (Fig. 5j). The magnitude of this difference across latent coordinates and time, signed by the change in the C5 trial factor, gave one inhibition-related change per session and inhibition type (Methods). The signed C5 change showed a modest association with the matching accuracy change (*ρ* = 0.266, *n* = 180 session–inhibition observations; Fig. 5k), with larger shifts in C5 accompanying poorer performance. Components selected for instruction, choice and outcome showed weaker, non-significant associations (Supplementary Fig. S5; Table S2). These analyses demonstrate Neuralign’s capacity to compare perturbation responses across heterogeneous brain-wide recordings, with a single shared component linking inhibition-related changes in neural activity to differences in behavioural susceptibility across recordings.

### 2.6 Inhibition remains detectable outside the choice direction on error trials

Having identified a shared inhibition response, we next asked whether it was confined to the dominant choice direction, and whether it remained detectable on error trials. Earlier work has linked choice errors to changes in ALM selectivity [30], and models suggest that activity outside task-coding directions can strongly influence decision trajectories [31]. To address this across all 93 sessions, we used the aligned projection of measured activity *Q*^(*n*)⊺^*Z*^(*n*)^, rather than the fitted components; because this projection is linear and fixed once fitted, a reference learned only from uninhibited, correct trials could be applied to reserved inhibition and error trials.

We fitted a separate rank-10 condition-aware Neuralign model, anchored by left and right choices, to a balanced subset of correct, uninhibited trials from each session, reserving all remaining trials for evaluation (Methods; rank selection in Supplementary Fig. S6e,f). We regressed out licking and tracked jaw, nose and tongue movements from the aligned neural activity, fitting the regressions on the training subset only. The choice direction was defined as the leading direction of the right-correct minus left-correct difference in mean activity during the final half-second of the delay, which captured more than 95% of this difference in every fit, and its orthogonal complement defined a nine-dimensional choice-residual subspace (Fig. 6a).

**Fig. 6.**
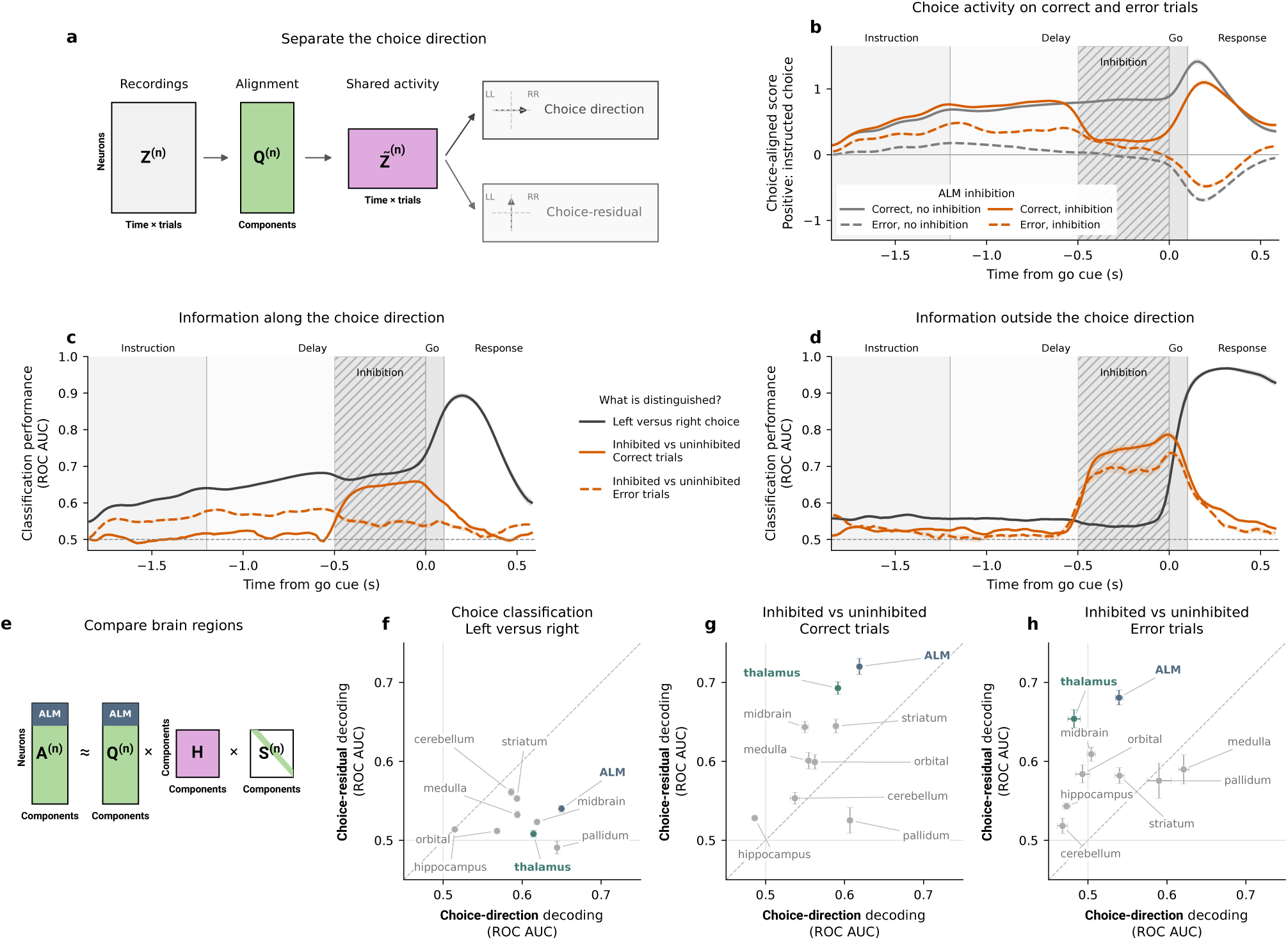
Inhibition remains detectable outside the choice direction on errors. **a**, Shared activity is projected along the choice direction or into its orthogonal, choice-residual subspace. The reference uses correct trials without inhibition; LL and RR denote correct left (left instruction + left choice) and correct right trials. **b**, Choice-aligned activity for correct (solid) and error (dashed) trials, without inhibition (grey) or with bilateral inhibition (orange). Positive scores point towards the instructed choice. Negative scores indicate activity moving in the uninstructed (error) direction. **c,d**, Held-out-session temporally resolved decoding along (**c**) or outside (**d**) the choice direction. Charcoal denotes choice decoding and orange inhibition decoding on correct (solid) and error (dashed) trials; the horizontal dashed line marks chance ROC AUC. **e**, Matching rows of the neuron factors and *Q* identify regional contributions; for example, ALM rows are highlighted. **f–h**, Regional choice decoding (**f**) and inhibition decoding on correct (**g**) and error (**h**) trials, along the choice direction versus the choice-residual subspace during the inhibition period; the diagonal marks equal performance. ALM and thalamus are highlighted. Lines and shading in **b–d**, and bars in **f–h**, show mean *±* s.e.m. across ten fits and training-trial splits.

Along the choice direction, activity on error trials pointed less strongly towards the instructed choice than on correct trials throughout the delay (Fig. 6b). On correct trials, bilateral inhibition shifted activity towards the incorrect choice along the choice direction during inhibition, and activity recovered after inhibition ended. We then trained time-resolved classifiers, evaluated on held-out sessions, to distinguish left from right choices within each instruction, and bilateral inhibition from no inhibition separately on correct and error trials (Fig. 6c,d). As expected from its construction, choice decoding during the late delay was stronger along the choice direction than outside it.

In contrast, inhibited and uninhibited trials were more distinguishable outside the choice direction than along it, on both correct and error trials, and this residual-subspace signal rose sharply at inhibition onset (Fig. 6c,d). On error trials, late-delay inhibition decoding reached an area under the receiver operating characteristic curve (AUC) of 0.685 ± 0.011 in the choice-residual subspace, compared with 0.548 ± 0.005 along the choice direction (mean ± s.e.m. across ten fits and training splits). Higher residual-subspace decoding was consistent across the ten fits (Supplementary Fig. S6c), and although the residual subspace has more dimensions than the choice direction, shuffled-label controls remained at chance in both, indicating that this difference is meaningful (Supplementary Fig. S6a,b). Thus, inhibition-related information persisted on error trials after removing the dominant choice direction and fitted movement effects.

Because the projection is linear and each row of *Q*^(*n*)^ corresponds to a recorded neuron, we resolved regional contributions to choice and inhibition decoding during the inhibition period by recomputing the projection with each region’s neurons separately (Fig. 6e). Choice decoding was generally stronger along the choice direction (Fig. 6f). In contrast, on correct trials, inhibition was more detectable outside that direction than along it in every region except pallidum, most strongly in the ALM, thalamus, striatum and midbrain (Fig. 6g). On error trials, the ALM and thalamus retained the largest choice-residual inhibition scores, and in thalamus inhibition was detectable only outside the choice direction (Fig. 6h).

ALM inhibition therefore left a signal outside the dominant choice direction, including on error trials, with the strongest contributions from ALM and thalamus. This dissociation is consistent with earlier ALM recordings in which choice-coding activity recovered after unilateral inhibition whilst a separate, non-choice-selective mode retained the perturbation signature [32], and with the observation that ALM inhibition alters activity and choice selectivity in connected subcortical regions [29]. Neuralign extends these observations across 93 sessions with different regional sampling and resolves where distributed perturbation signals remain detectable outside the dominant choice direction.

## 3 Discussion

Population activity patterns recur across days and animals performing the same task [5, 6]; however, their distribution across neurons changes between sessions and their strength is expressed variably across trials. There is growing recognition that such activity patterns are well described by a small number of components, each with a characteristic time course [20]. However, identifying these components across recordings or sessions is complicated by the inability to accurately align and decompose such data. Linear alignment methods such as canonical correlation analysis, SRM and Hyperalignment return a common space in which recordings can be compared [9, 10], but do not explicitly factor each shared motif into temporal, trial and neuronal contributions. With Neuralign, we have shown that recordings containing different neurons and variable numbers of trials can be represented as a joint decomposition into components with shared temporal factors and session-specific trial and neuron factors. This structure makes the representation directly interpretable: each neural motif can be followed through its timing, its strength across trials and its contribution to different populations. We therefore believe that viewing large-scale modern neuroscience datasets through the lens of the Neuralign decomposition offers a productive route for linking biological questions to preserved dynamical motifs.

In simulations, recovery of the generating factors improved as we incorporated more sessions, most markedly from two to twelve sessions, significantly exceeding that of TCA fitted to each session separately. The learned session gains also retained the grouping of recordings. In recurrent networks, trial-factor distributions distinguished related from distinct dynamical solutions. Neuralign’s shared representation can therefore improve recovery of common structure whilst preserving differences between the systems generating the activity. In experimental recordings, we showed that Neuralign factors supplied interpretable readouts. Fitted to motor-cortex recordings, without reach or region labels, Neuralign recovered the ring-like organisation of reach directions in the trial factors and regional differences between M1 and PMd in the neuron factors, whilst session gains from the label-free fit grouped recordings by animal.

Using the Mesoscale Activity Map dataset comprising 93 brain-wide recordings, we managed to isolate the shared inhibition response in a single component using an unsupervised Neuralign fit, validated in held-out sessions. This single component distinguished ALM inhibition conditions across recordings with different neuronal sampling, and its change showed a modest association with behavioural impairment. In contrast, SRM and Hyperalignment do not isolate the shared inhibition signal into a single component.

The linear projection of a condition-aware reference, learned from correct, uninhibited trials, extended the analysis to trials never used for fitting, revealing inhibition-related activity outside the dominant choice direction on both correct and error trials. Whereas the contribution of ALM provides a positive control, the thalamic contribution, detectable only outside the choice direction on error trials, is consistent with the thalamocortical loop that maintains ALM preparatory activity [33], and with perturbation signals that persist outside choice-coding activity [32]. Neuralign makes that information comparable across recordings and traceable to individual trials and regions.

Neuralign is complementary to non-linear and multi-session models, including LFADS and related dynamical models, CEBRA, MARBLE and decoding-oriented alignment [12–16, 19], which can capture richer structure. For example, in our benchmarks, CEBRA transferred reach-target decoding across animals more effectively (Supplementary Fig. S3a). These models offer other forms of interpretability, such as the flow geometry described by MARBLE [16], but not explicit factors for each neuron, trial and recording. Within animals, hand-position decoding from Neuralign improved on SRM and Hyperalignment, although Neuralign was evaluated on rank-constrained, factor-predicted trajectories, so part of this gain may reflect de-noising; interpretable structure therefore need not come at the expense of behavioural readout within a cohort. In practice, Neuralign is best suited to questions about which pattern is shared, when it occurs, on which trials and in which neural populations.

Several limitations accompany these conclusions. First, component interpretation requires stability across fits; conditional uniqueness results for tensor decompositions [20] do not establish identifiability of the complete coupled model. Second, in label-free fits, the signs of a component’s session gain and trial factors can be exchanged within a recording without changing the loss, so signed comparisons across recordings depend on an explicit convention. Third, decoder transfer was evaluated within jointly fitted cohorts and did not significantly improve across animals, so aligning new recordings to a fixed reference remains an important next test, with direct relevance to stabilising brain–computer interfaces [11]. Neuralign also assumes a common time axis and shared temporal profiles, and its trial factors capture changes in amplitude rather than timing; because our simulations were generated under these assumptions, robustness to latency variability remains to be tested, and time-warping approaches [34] offer a natural extension. Finally, session-specific activity is accommodated through the soft coupling rather than modelled explicitly, and the aligned projection is linear, so explicit separation of shared and private processes [35] would be a useful extension.

Neuralign provides an interpretable shared representation in which a neural motif remains linked to the trials and populations expressing it. As recordings scale to brain-wide, chronic and multi-animal datasets [5, 6, 29], the central question shifts from whether neural activity is shared to how shared activity is expressed across trials, regions, individuals and perturbations. Neuralign addresses this question directly, with immediate applications in comparing perturbation responses across cohorts with heterogeneous regional sampling and in following component expression through learning in chronic recordings, and, once new recordings can be aligned to a fixed reference, in relating each new session to an established set of shared components. Its broader value is to make shared neural activity accessible to biological questions: how conserved motifs vary with movement and behavioural outcome, and how their expression changes across neural populations.

## 4 Methods

### 4.1 Neuralign model

Neuralign models *N* neural data tensors 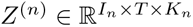, with neuron, time and trial modes. Neuron and trial counts may differ across sessions, but the time axis is shared. The model combines a canonical polyadic (CP) reconstruction [36] with session-varying factors inspired by PARAFAC2 [22]:

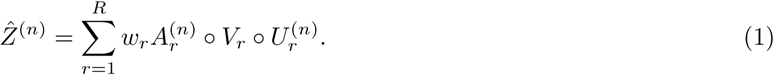

Here *R* is the number of components, ◦ denotes the outer product, 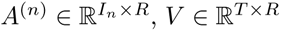 and 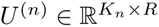 are neuron, temporal and trial factors, respectively, and *w* ∈ ℝ*^R^* contains global component weights.

The model also constructs aligned neuron factors

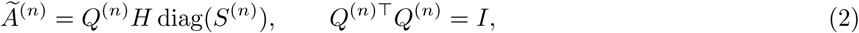

where 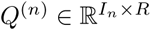 is a session projection, *H* ∈ ℝ*^R×R^* is the shared mixing matrix and *S*^(*n*)^ ∈ ℝ*^R^* contains session gains. The soft coupling between *A*^(*n*)^ and *Ã*^(*n*)^ follows flexible PARAFAC2 [23]. Trialor condition-averaged trajectories orient the session projections, while reconstruction retains individual trials.

For label-free alignment, let 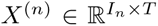 be activity averaged over the trials used for alignment and *u̅*^(^*^n^*^)^ their mean trial-factor row. The model-predicted mean trajectory is

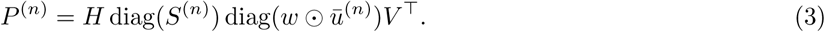

The projection is updated by orthogonal Procrustes:

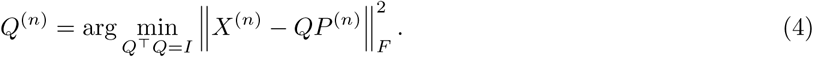

This update is computed from the singular-value decomposition of *X*^(*n*)^*P* ^(*n*)^*^⊤^*. With the projections fixed, the differentiable objective combines normalised reconstruction and neuron-factor alignment (coordinate) errors:

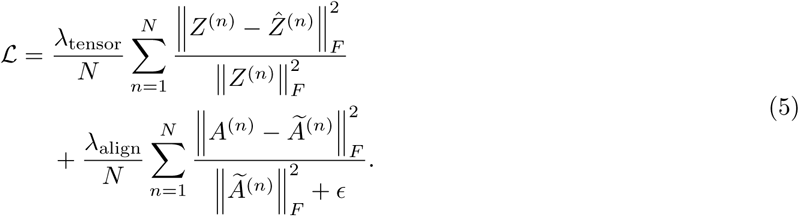

Both loss weights were set to one, with *ɛ* = 10*^−^*^12^. Reconstruction errors are normalised by tensor energy, and both terms average sessions equally. The mean-trajectory projection update is performed separately from gradient optimisation of this objective.

For condition-aware alignment, trial factors are averaged within each condition and then across sessions to obtain shared condition prototypes *C_c_*. Condition-averaged neural activity and its predicted trajectory

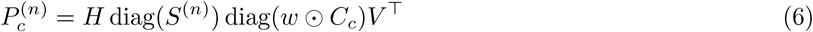

are stacked across labels before solving the same Procrustes problem. Labels orient the projections but do not enter the reconstruction term.

Factors were initialised from leading singular vectors of concatenated trial means or their pooled temporal covariance, with trial factors initialised by least squares. Fits used signed, unconstrained factors and the Adam optimizer with a learning rate of 10*^−^*^2^; projections were recomputed every three steps without differentiating through the Procrustes update. Temporal and mixing columns were normalised to unit Euclidean norm, and trial-factor columns to unit root-mean-square magnitude with equal session weighting. Their scales were absorbed into *w* or *S*^(*n*)^, preserving reconstruction and aligned predictions. Early stopping used a tolerance of 10*^−^*^6^ and patience of 100 steps.

### 4.2 Synthetic factor-recovery benchmark

Each simulated dataset contained two shared components, with 64 time points, 8–32 neurons and 8–32 trials per session. Shared temporal factors were a sigmoid and gamma-shaped transient; their trial factors decreased or increased across trials. The shared mixing matrix had diagonal entries of one and off-diagonal entries of 0.1 before column normalisation. Session projections were orthonormalised from synthetic neuron loadings with either one or two spatial groups (regions). Session gains came from two centres, (1, 0.1) and (0.1, 1). Each session also contained one private component with gain 0.01–0.1. The resulting rates had a nominal 10-Hz baseline and 5-Hz modulation scale; neuron baselines were raised when needed to keep rates at least 1 Hz. Spike counts were sampled independently from these rates in 50-ms bins. All methods received the same spike counts after subtraction of the known baseline. No temporal smoothing or neuron-wise z-scoring was applied.

Recovery was evaluated with rank-2 Neuralign, SRM and Hyperalignment, matching the two shared components. TCA was fitted separately to each session at rank 3 to include the private component. SRM and Hyperalignment were fitted to trial averages and then applied to individual trials. Cohorts contained 2, 12, 24 or 36 sessions, with 10 simulated datasets per cohort size. Factor columns were matched to ground truth indepen-dently within each mode by maximising absolute cosine similarity (factor match score), allowing for component order and sign. Scores were averaged across the two shared components and sessions for each mode, then across the neuron, temporal and trial modes for the overall score. SRM and Hyperalignment supplied a shared temporal response as the temporal-factor proxy, session projection columns as the neuron-factor proxy, and time-averaged single-trial embeddings as the trial-factor proxy. Confidence intervals used bootstrap resampling of the 10 simulated datasets. Supplementary rank controls used a separate fixed 12-session Poisson dataset, fitted at ranks 1–8 with three initialisation seeds per rank.

### 4.3 Recurrent networks and preprocessing

The RNN comparison used three pre-trained 500-unit, rank-2 recurrent-network checkpoints and their precomputed delayed match-to-sample trajectories from the MARBLE analysis [16], following the low-rank network construction of Dubreuil, Valente et al. [27]. The trajectory analysis compared 20 input gains linearly spaced from 1 to 0. The retained series contained 100 trials per gain for I/A and I/B and 200 for II.

Original trials comprised fixation, sample, delay, test and response epochs lasting 0.5, 0.5, 3, 0.5 and 1 s, respectively, at 20-ms resolution. The first 15 steps of each epoch were removed, leaving a shared 200-bin axis. For each network, 200 units were selected without replacement using a fixed random generator, and each unit was z-scored across retained time bins and trials. Unconstrained, label-free Neuralign models were fitted at ranks 1–8 with three initialisations per rank. Rank 5 was selected by inspecting reconstruction and alignment errors (Supplementary).

### 4.4 RNN projections and trial geometry

Measured trajectories were obtained as *Q*^(*n*)^*^⊤^Z*^(*n*)^. Displayed trajectories used 16 trials per gain and the first three aligned coordinates. Session-gain plots show the first three signed entries of *S*^(*n*)^.

Pairwise Gaussian 2-Wasserstein distances quantified dissimilarity of trial factors across the 60 network–gain groups. Metric multidimensional scaling embedded this distance matrix in two dimensions.

### 4.5 Macaque reaching data and preprocessing

The analysis used the Perich–Miller centre-out reaching data from Dandiset 000688 [28, 37]. Spikes were aligned to the go cue and counted in 20-ms bins from −0.5 to 0.8 s. Sessions were retained if they included both PMd and M1, at least 10 units per region and at least 10 trials per reach target. Units with no spikes were removed, leaving 36 sessions (14 from monkey C, 20 from M and 2 from T). Each neuron was z-scored across time and trials and smoothed in time with a Gaussian kernel (50-ms standard deviation).

For the factor analysis, label-free models of rank 1–16 were fitted with three seeds. Rank 6 was selected by inspecting reconstruction and alignment-error curves (Supplementary Fig. S3e,f). Reach target was decoded from *U* ^(*n*)^ and region from *A*^(*n*)^ using LDA (linear discriminant analysis) with five-fold stratified cross-validation. Standardisation used training data only. Permutation importance was the decrease in balanced accuracy after shuffling one component in the test fold (10 repeats). Importances were averaged over folds within each session and then equally across sessions. Animal identity was decoded from the six session gains of the same label-free fit, using training-only standardisation and two stratified session folds. Ten held-out permutations per component identified the two gains plotted directly in Fig. 4j. The supplementary principal component analysis (PCA) used a separate condition-aware fit after standardising each gain across sessions.

### 4.6 Joint-fit decoder transfer

Rank-6 Neuralign, SRM, Hyperalignment and CEBRA representations were fitted jointly to all 36 sessions in label-free and condition-aware modes. Each mode used ten fits; condition-aware alignment used the eight reach identities. SRM and Hyperalignment were fitted to trial-averaged trajectories in label-free mode and concatenated target-averaged trajectories in condition-aware mode, then applied to individual trials. SRM was implemented using fastsrm, with sessions zero-padded to the largest neuron count. Hyperalignment used rank-6 PCA for each session followed by generalised Procrustes alignment in nltools. Multi-session CEBRA used a shared time index in label-free mode and time plus one-hot reach identity in condition-aware mode. CEBRA used the offset10-model architecture, six output dimensions, cosine distance, time-delta conditioning with an offset of ten bins, a learning rate of 3×10*^−^*^4^, batch size 512 and 500 iterations. Neuralign was evaluated using reconstructed latent trajectories.

Decoders used five condition-stratified folds per fit. All sessions contributed to representation fitting, including those held out from decoder training. Within-session decoders used training trials from the target session; within-animal decoders used the other sessions from that animal; and leave-one-animal-out decoders used sessions from the other two animals. For target decoding, flattened trial trajectories were reduced to six PCA features and classified by LDA (non-Neuralign methods). PCA used decoder-training trials only. For kinematics, trials were split before post-go time bins were flattened; predictors and two-dimensional position or velocity targets were standardised on decoder-training data and fitted by ridge regression. Performance was measured by balanced accuracy for targets and *R*^2^, averaged across the two coordinates, for kinematics. Reported cohort means weight sessions equally after averaging decoder folds and representation repeats within each session.

### 4.7 Mesoscale Activity Map cohort

The analysis used the Mesoscale Activity Map delayed-lick recordings in Dandiset 000363 [29, 38]. Trials with early licks, automatic water or free water were excluded; hit and miss trials were retained. Units labelled “good” by the source dataset were grouped into ALM, orbital cortex, striatum, pallidum, hippocampus, thalamus, midbrain, medulla and cerebellum. Sessions required ALM, at least 50 retained units, at least 50 correct trials for each instruction and at least 10 photostimulation trials. Only late-delay photostimulation trials (−0.5 to 0 s relative to the go cue) were included. A total of 93 sessions from 19 mice met these criteria.

Spikes were aligned to the go cue and counted in 20-ms bins from −1.85 to 0.6 s. For the component analysis, spikes were smoothed with a Gaussian kernel (50-ms standard deviation) and z-scored per neuron across time and trials. Label-free models at ranks 1–16 were fitted with three seeds (Supplementary Fig. S4d,e). The component analysis used the rank-12 fit with the lowest reconstruction error. The error analysis used rank 10 and left/right labels from correct, uninhibited trials, with ten seeds and training-trial splits. Its normalisation used training trials only, as described below; the condition-aware rank sweep is shown in Supplementary Fig. S6e,f.

### 4.8 Behavioural impairment, component decoding and latent reconstruction

The probability of a rightward choice was modelled by Bayesian mixed-effects logistic regression. Fixed effects were instruction, inhibition type, their interaction and animal identity. Random effects included a session intercept and animal-specific slopes for the instruction-by-inhibition interactions. Fitting used the variational Bayes implementation in statsmodels, with zero-mean Gaussian priors of standard deviation 2 for fixed effects and 0.5 for log random-effect standard deviations. Five-fold cross-validation was stratified jointly by session and photostimulation presence. For each held-out trial, posterior-mean coefficients, including random effects, gave correct-choice probabilities under no, unilateral and bilateral inhibition. Session impairment was the predicted accuracy difference averaged within each fold and then across folds, expressed as a fraction. Negative values indicate poorer performance without trial-level inhibition information.

Component importance was evaluated separately for instruction (left/right), choice (left/right), outcome (hit/miss) and inhibition (none/unilateral/bilateral). Instruction, choice and outcome used uninhibited trials; inhibition used all trials. Within each session, LDA with training-only standardisation used five stratified folds, reduced to the smallest class count when necessary. Sessions missing a required class or containing fewer than two trials in a class were excluded for that feature. Each component was permuted ten times on held-out trials, and importance was the decrease in balanced accuracy. Fold means were averaged with equal session weights. The component with the largest mean importance was selected for each feature, with ties resolved by component number.

Decoder transfer used ten-fold stratified group cross-validation with session as the group. For each outer fold, the component was selected by averaging only the training sessions’ within-session importances; scaling and LDA were then fitted to training trials and evaluated on held-out sessions. The decomposition included all 93 sessions. For comparison, SRM and Hyperalignment were each fitted once at rank 12 to label-free trial averages from the same preprocessed recordings, using the implementations above (seed 42). Full-trial time means of native aligned coordinates served as trial-factor proxies, with the same importance procedure and identical session folds used to select and evaluate one coordinate. These readouts differ from Neuralign’s learned temporal-factor amplitudes and test native-coordinate interpretability rather than decoding from the full aligned subspace.

For each component selected across all sessions, we reconstructed its contribution to the Neuralign decomposition in aligned coordinates. Inhibition-related change was the Frobenius norm of the difference between inhibited and uninhibited trial means, signed by the inhibited-minus-uninhibited change in its mean trial factor. Each change was paired with the corresponding model-estimated accuracy change, yielding 180 session–inhibition observations.

### 4.9 Choice-subspace and regional error analysis

For each of ten seeds, equal numbers of correct left and correct right uninhibited trials were selected per session, taking 80% of the smaller class count for each side (rounded down). These training trials determined neural and movement normalisation, the rank-10 condition-aware fit and the movement regressions. The remaining correct trials, spontaneous errors and inhibited trials were reserved for evaluation.

Movement regressors were licking and the *x* and *y* coordinates of the jaw, nose and tongue, standardised using training trials. One ridge regression (penalty one, no intercept), pooled across training trials from all sessions and time bins, predicted aligned activity from movement. Its predictions were subtracted from all trials. The choice direction was obtained by singular-value decomposition of the session-averaged right-correct minus left-correct training trajectory from −0.5 to 0 s. The smallest number of singular vectors explaining at least 95% of the squared contrast was selected; this was one in every fit. Its orthogonal complement defined the nine-dimensional choice-residual subspace. For the choice-aligned score, projections were centred at the midpoint of the correct left/right training means and divided by half their late-delay separation. The sign was then aligned to the instruction, so positive values point towards the instructed choice.

At each time bin, logistic regression with training-only standardisation decoded three contrasts: left versus right choice, bilateral inhibition versus no inhibition on correct trials, and bilateral inhibition versus no inhibition on error trials. Decoders were fitted separately for each instruction and their scores averaged; within a fixed instruction, the choice contrast also distinguishes correct from error trials. Five-fold stratified group cross-validation held out sessions from decoder training; all sessions contributed correct, uninhibited training trials to the reference and movement regressions. Performance was measured by area under the receiver operating characteristic curve (ROC AUC). To localise regional contributions, *Q*^(*n*)^*^⊤^Z*^(*n*)^ was recomputed using only neurons in one region. Each regional movement regression was fitted using training trials from sessions containing that region. Sessions lacking a region were excluded from its regional analysis. Choice-direction estimation and regional AUC summaries used −0.5 ≤ *t* ≤ 0 s.

### 4.10 Statistics and reproducibility

All sessions meeting stated inclusion criteria were analysed. Repeated Neuralign fits quantified variability across initialisations and, for the error analysis, training-trial splits.

Synthetic factor recovery at 12 versus 2 sessions was compared across paired data seeds using a one-sided Wilcoxon signed-rank test. Motor decoder comparisons used one-sided paired Wilcoxon signed-rank tests across the 36 target sessions, testing whether Neuralign exceeded each baseline. Session scores were averaged over decoder folds and representation repeats before testing. Benjamini–Hochberg correction used a 24-comparison family for Fig. 4 (four evaluation settings, three outcomes and two baselines). Supplementary motor tests used a 54-comparison family (two representation modes, three evaluation settings, three outcomes and three baselines).

Component–behaviour associations used Spearman correlation. Two-sided *p*-values were computed from 1,000 label permutations with a plus-one correction; inhibition-specific impairment was permuted within inhibition type. Component–behaviour *p*-values were unadjusted across features. Permutation-test annotations use \**p <* 0.05, \*\**p <* 0.01 and \*\*\**p <* 0.001; ns denotes not significant. Decoder annotations use the corresponding thresholds for BH-adjusted *q*-values. Analyses were not blinded.

## Supplementary information

Supplementary figures provide factor-recovery and rank checks (S1–S2), motor transfer benchmarks (S3), cohort and behavioural controls (S4), task-feature components and alignment controls (S5), and choice-subspace controls (S6). Numerical source tables accompany the analysis code. Repeated fits measure initialisation or training-split variability rather than biological replication.

**Fig. S1.**
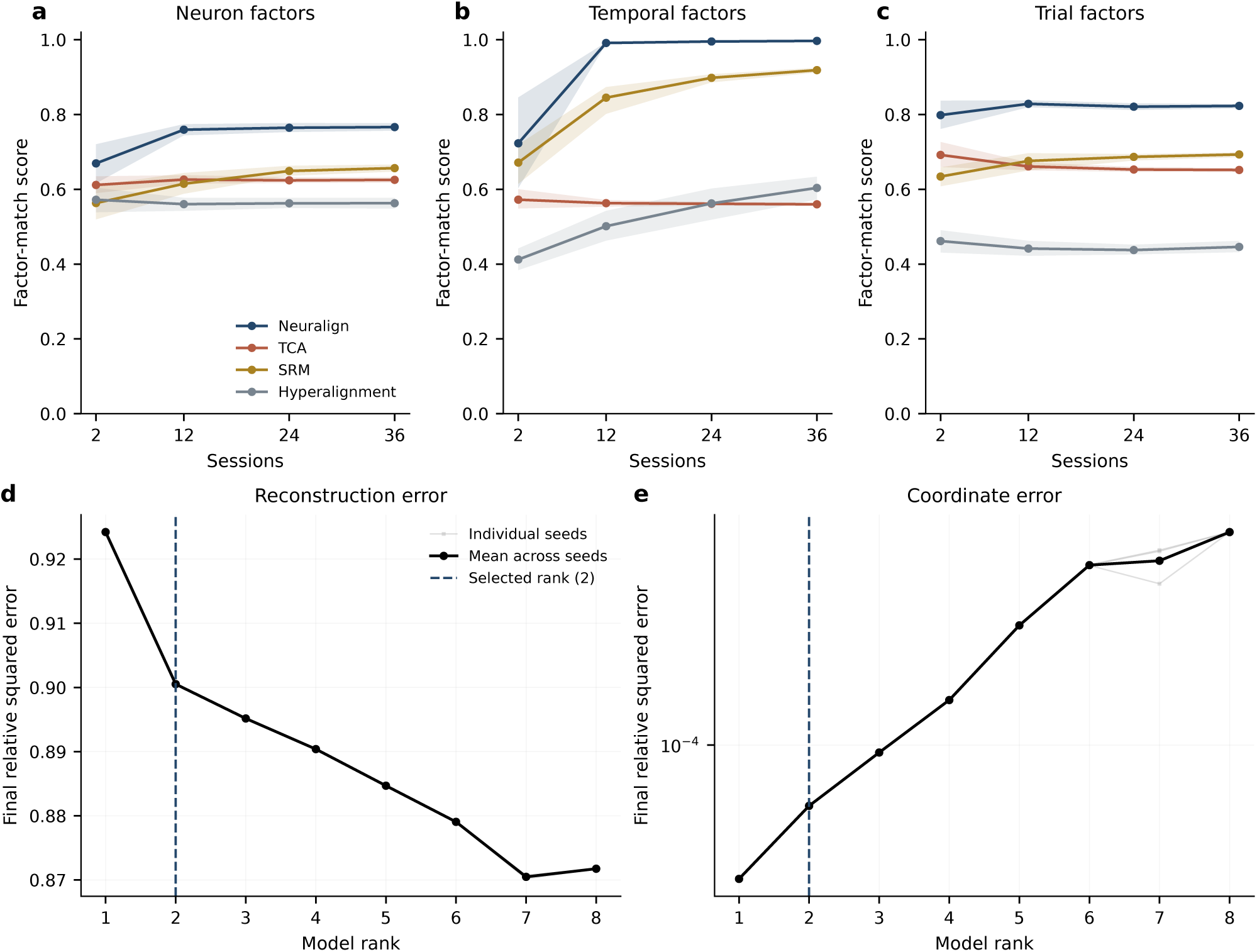
Synthetic factor recovery and rank controls. **a–c**, Neuron, temporal and trial factor recovery across ten simulated datasets, using the same data as Fig. 2. Lines show means and shading bootstrap 95% confidence intervals. **d,e**, Final reconstruction and alignment errors for ranks 1–8 on a fixed 12-session dataset. Thin grey lines show three initialisations and the black line their mean; the dashed line marks the known shared rank of two.

**Fig. S2.**
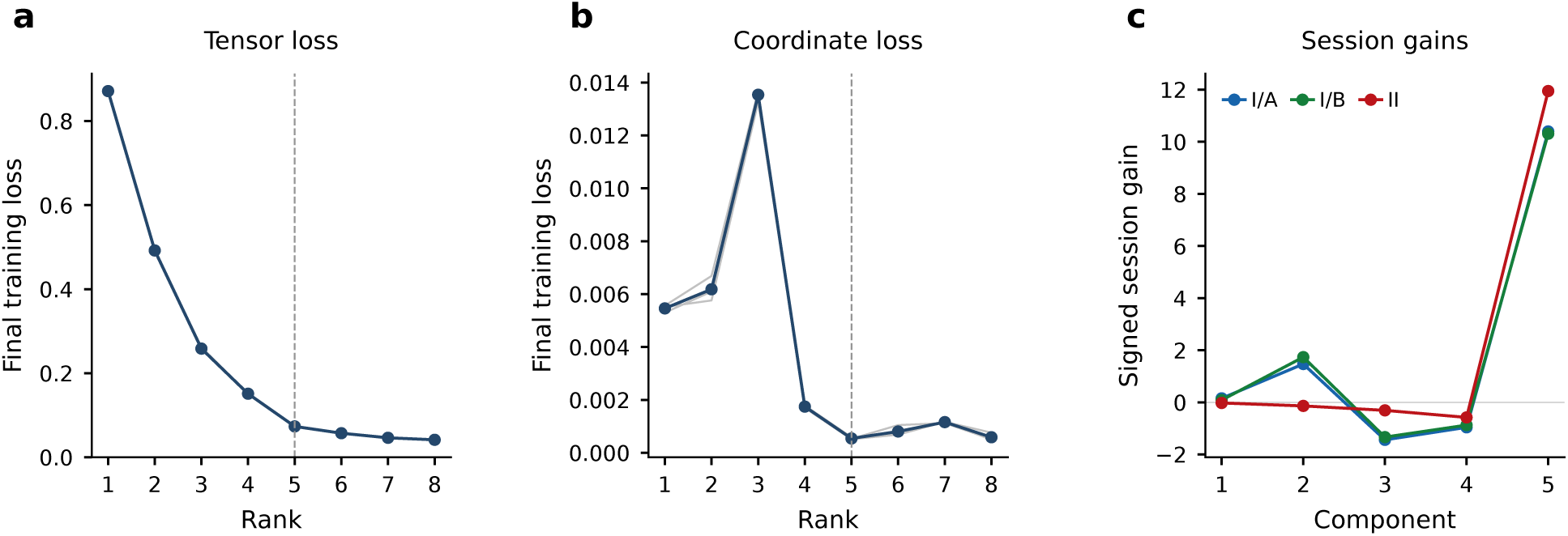
RNN rank selection and session gains. **a,b**, Final reconstruction and alignment losses for ranks 1–8. Thin grey lines show three initialisations and the navy line their mean; the dashed line marks the selected rank of five. **c**, Signed session gains across all five components for I/A (blue), I/B (green) and II (red), using the same fit as Fig. 3.

**Fig. S3.**
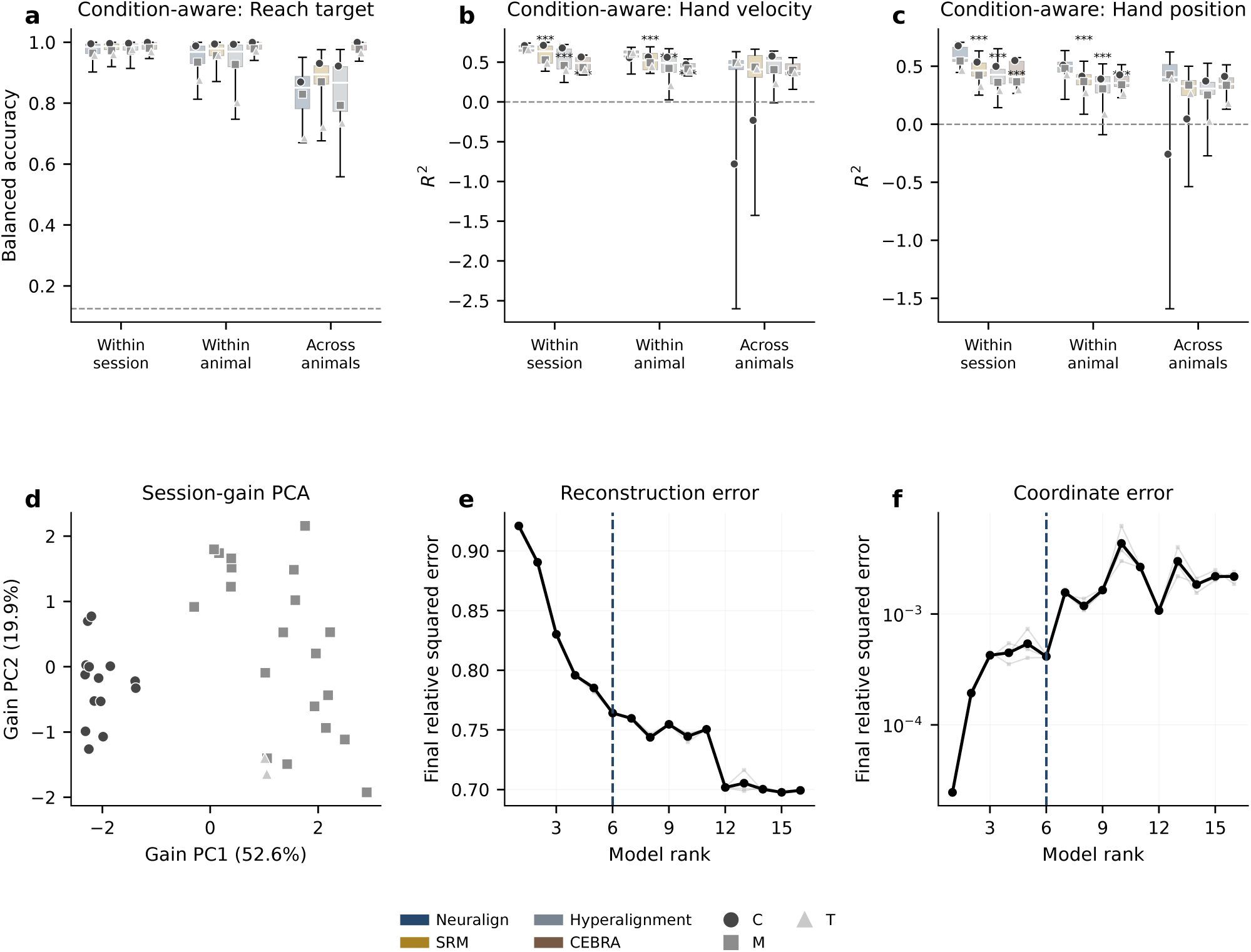
Motor decoder transfer and session gains. **a–c**, Condition-aware decoding of reach target, velocity and position within sessions, across sessions within an animal and across animals. Boxes show medians and interquartile ranges across 36 sessions; whiskers span the full range, including negative *R*^2^. Grey markers show animal means (C, circles; M, squares; T, triangles). Dashed lines mark chance or zero *R*^2^. Asterisks denote BH-corrected one-sided paired Wilcoxon tests (\**q <* 0.05, \*\**q <* 0.01, \*\*\**q <* 0.001; 54-comparison family). Representations are jointly fitted; transfer refers to decoder training. **d**, PCA of standardised session gains, coloured and marked by animal; PC1 and PC2 explain 52.6% and 19.9% of variance. **e,f**, Rank sweeps showing three initialisations (grey), their mean (black) and selected rank 6 (dashed).

**Fig. S4.**
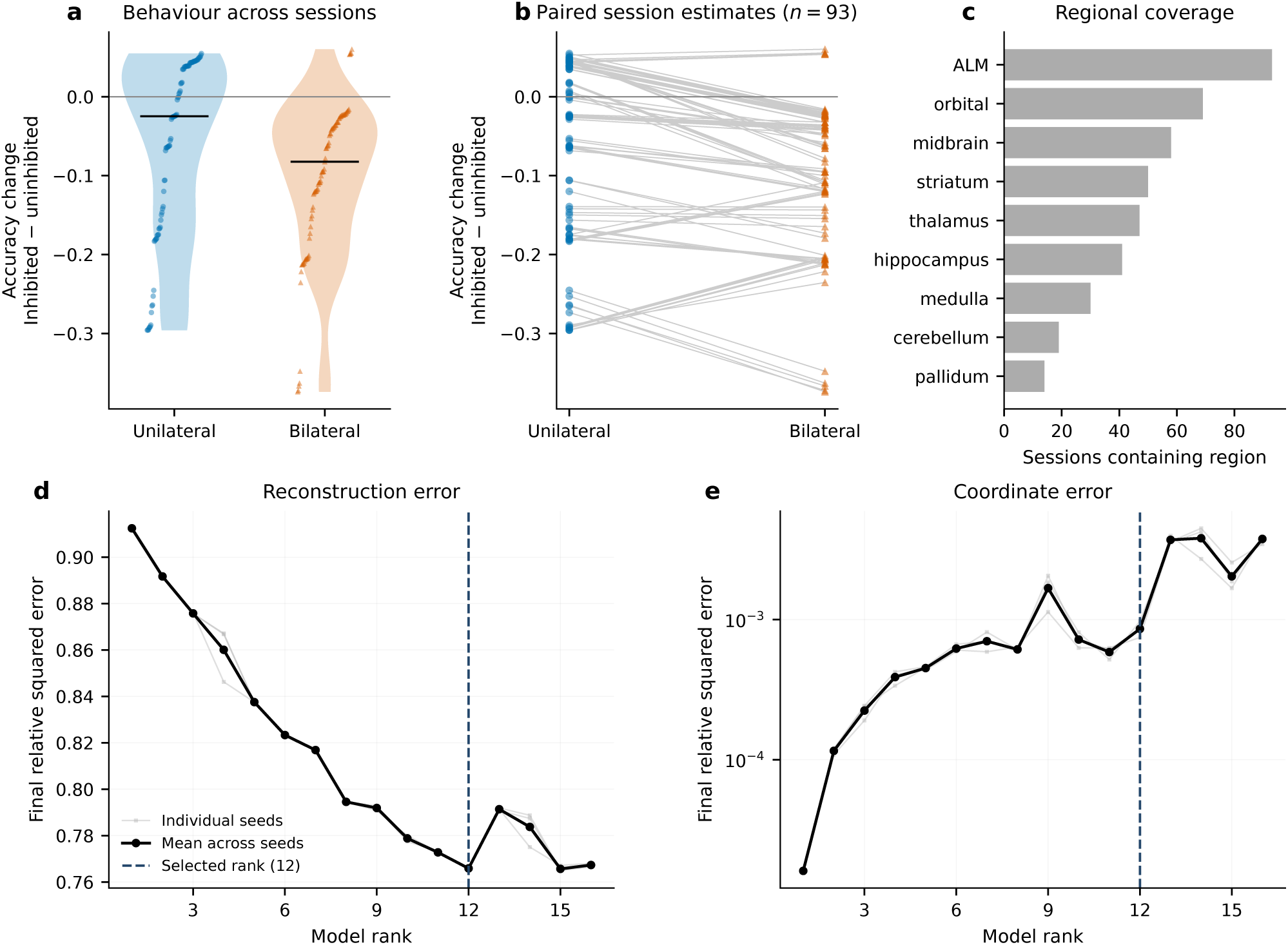
Behavioural susceptibility and regional sampling in the MAP cohort. **a,b**, Accuracy changes under unilateral and bilateral ALM inhibition across 93 sessions; negative values indicate impairment. Horizontal lines in **a** mark medians and paired lines in **b** join the same session. **c**, Sessions contributing neurons from each region. **d,e**, Label-free rank sweeps, showing individual initialisations (grey), their mean (black) and selected rank 12 (dashed).

**Fig. S5.**
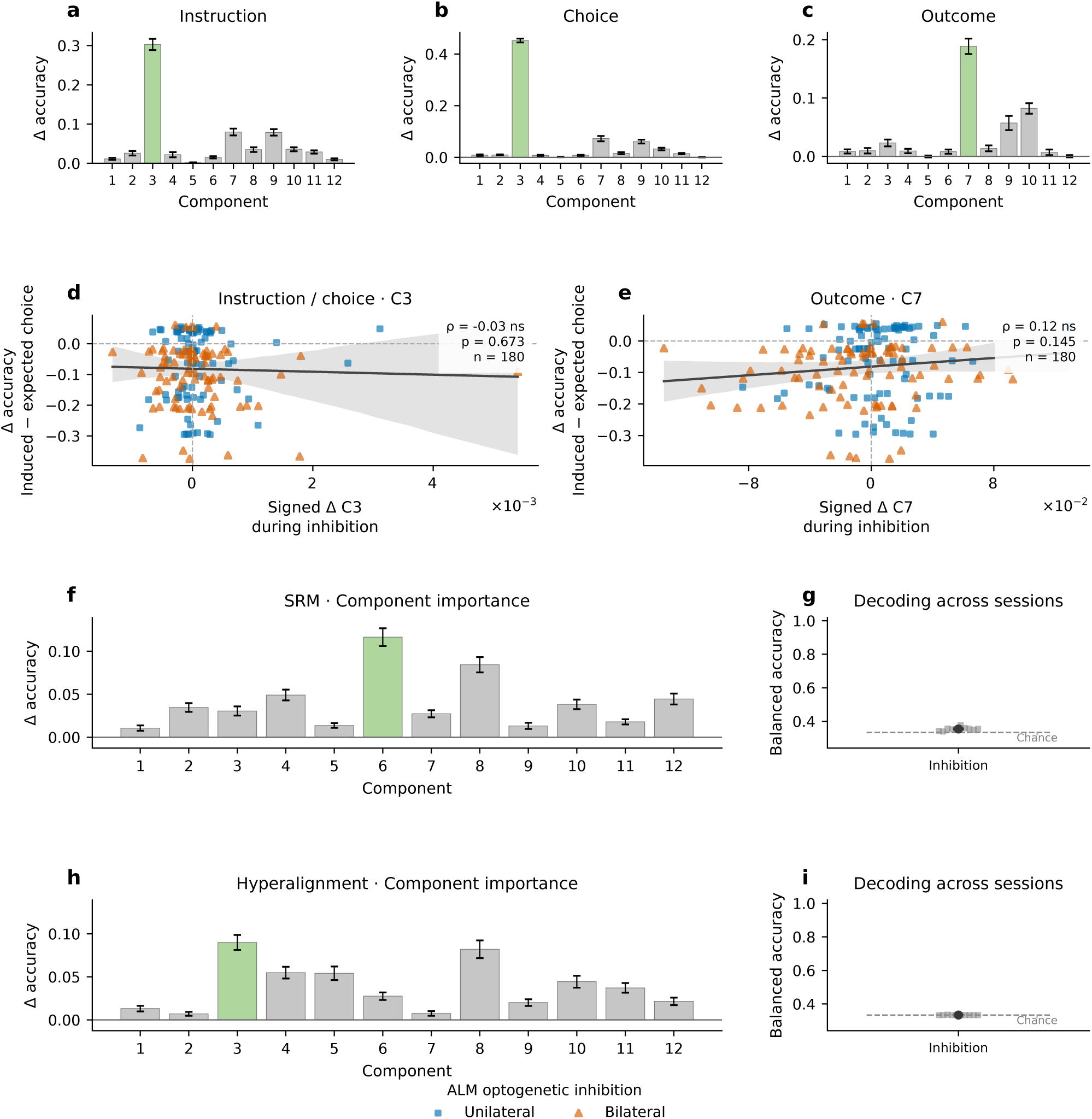
Task-feature components and single-component inhibition controls. **a–c**, Trial-factor importance for instruction, choice and outcome on uninhibited trials (mean *±* s.e.m. across 93 sessions); green marks the selected component. **d,e**, Signed late-delay latent changes against accuracy changes for unilateral (blue circles) and bilateral inhibition (orange triangles; 180 session–inhibition observations). Instruction and choice share component 3 (**d**); outcome selects component 7 (**e**). Reconstructions use unit global weights. Lines and shading show visualisation fits and session-bootstrap 95% confidence bands. Insets report Spearman correlations and unadjusted, two-sided permutation tests; ns denotes not significant. **f,h**, Inhibition importance across 12 native coordinates of label-free SRM (**f**) and Hyperalignment (**h**), using full-trial mean coordinate scores (mean *±* s.e.m. across 87 eligible sessions); green marks the selected coordinate. **g,i**, Inhibition decoding from one coordinate selected within each training fold. Small points show ten held-out-session folds; larger points and bars show mean *±* s.e.m.; dashed lines mark chance (1*/*3). All 93 sessions contribute to representation fitting.

**Fig. S6.**
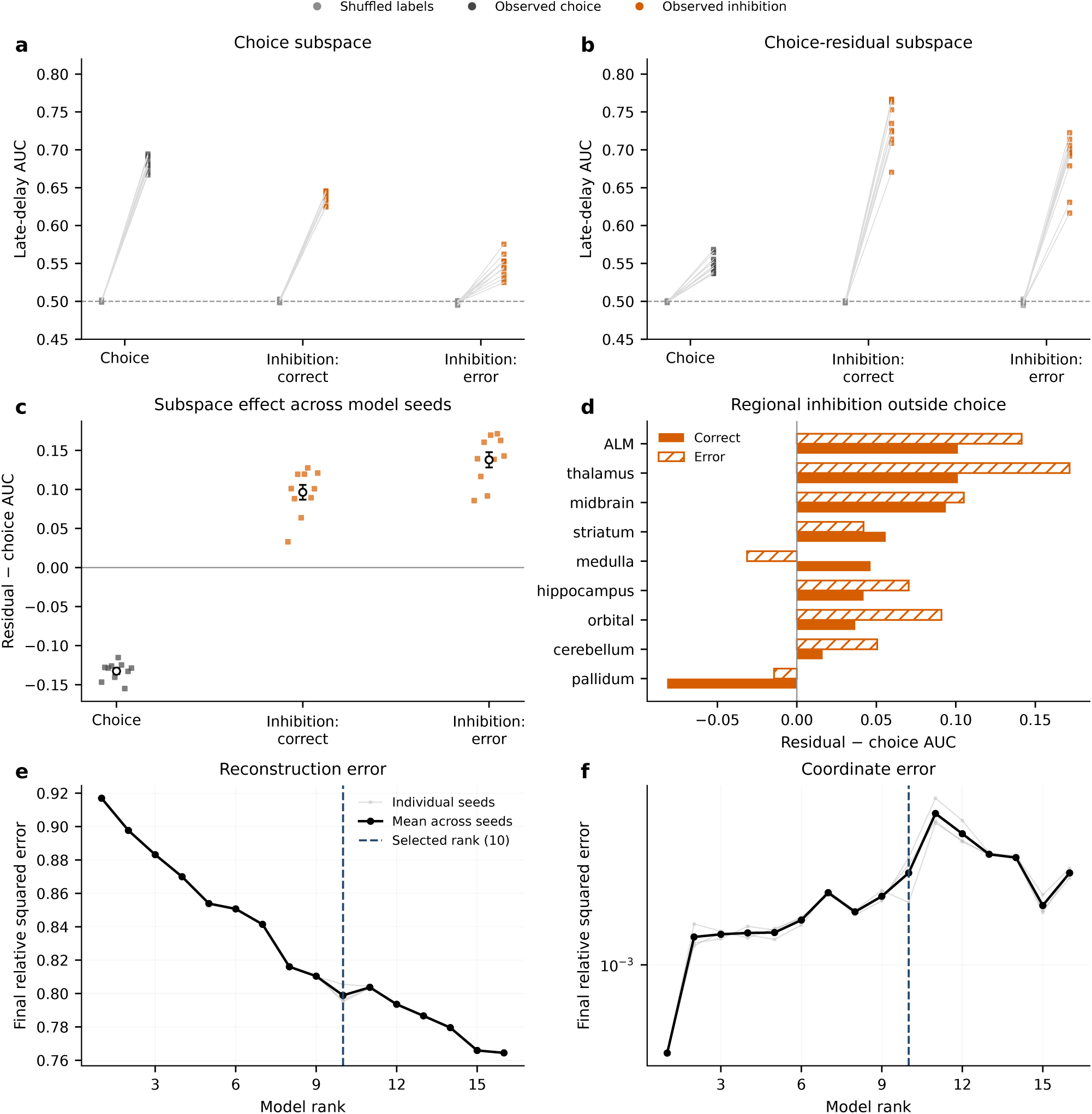
Controls for choice-subspace and regional decoding. **a,b**, Late-delay choice and inhibition decoding against shuffled-label controls. Joined points share the same fit and training split. **c,d**, Decoding in the nine-dimensional choice-residual subspace minus decoding along the one-dimensional choice direction, across task contrasts (**c**) and regions (**d**). Positive values favour the residual subspace. Small points in **c** show individual fits; open points and error bars show mean *±* s.e.m. across ten fits. Regional bars in **d** show differences of mean decoding scores. All decoding uses evaluation trials after movement regression. **e,f**, Condition-aware rank sweeps, showing individual initialisations (grey), their mean (black) and selected rank 10 (dashed). Alignment error uses a logarithmic axis.

**Table S1.** Datasets used in the main figures. The two ALM analyses use the same 93-session cohort, with separate fits. Synthetic datasets with the same seed share a prefix of sessions. The supplementary synthetic rank sweep uses a separate fixed 12-session dataset.

| Analysis | Sessions | Subjects or replicates | Neuralign rank |
|---|---|---|---|
| Synthetic (Fig. 2) | 2, 12, 24, 36 | 10 datasets per cohort size | 2 |
| RNN solutions (Fig. 3) | 3 | 3 networks | 5 |
| Reaching (Fig. 4) | 36 | 3 macaques | 6 |
| ALM inhibition (Fig. 5) | 93 | 19 mice | 12 |
| ALM error analysis (Fig. 6) | 93 | 19 mice | 10 |

**Table S2.**
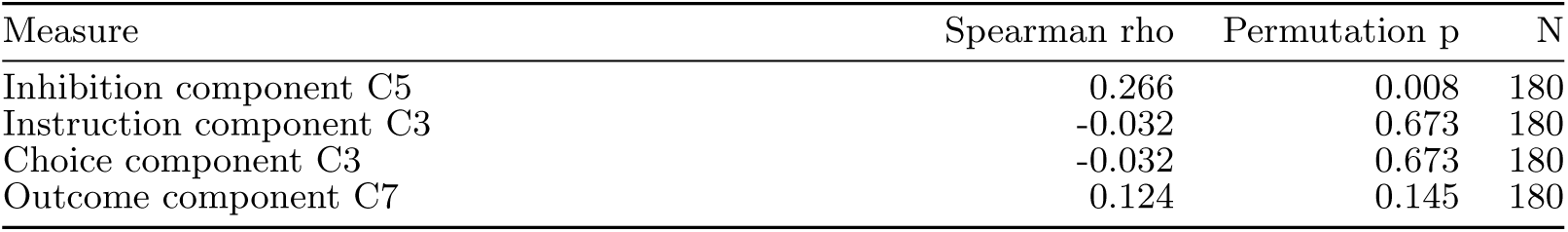
Associations between signed component changes and accuracy changes (inhibited minus uninhibited), using unit-weight latent reconstructions (Figs. 5 and S5). Instruction and choice use the same component. Two-sided permutation p-values use 1,000 within-inhibition-type shuffles and are unadjusted across features. N counts session–inhibition observations.

**Table S3.** Analysis and validation settings. Representation repeats are not biological replicates. Joint fits include decoder-test sessions; decoder transfer is conditional on that fitted representation. Figure 6 instead reserves evaluation trials from reference fitting and uses projected measured activity, not reconstructed trajectories.

| Analysis | Fit or data | Validation and readout |
|---|---|---|
| Synthetic recovery | 10 independent datasets per cohort size | Rank-2 shared-factor recovery; bootstrap intervals over datasets. |
| Motor factors | Best-loss label-free fit from 3 seeds | Five-fold LDA; training-only scaling; 10 held-out permutations per component; equal session weighting. |
| Motor decoding | 10 joint fits per representation mode | Five decoder folds; within session, within animal or across animals. Main figure: 24-test BH family; supplement: 54 tests. |
| Animal identity | Best-loss label-free motor fit | Two stratified session folds; LDA and 10 held-out permutations per component of the session gains. |
| ALM behaviour | Bayesian mixed-effects logistic model | Five held-out folds; fold-averaged accuracy with inhibition minus without inhibition (fraction; negative is worse). |
| ALM trial factors | Best-loss rank-12 fit from 3 seeds | Up to five within-session importance folds and 10 permutations; ten session-grouped transfer folds, selecting one component using training sessions only. |
| ALM latent change | Selected rank-1 latent contribution; global weights set to one | Spearman correlation and 1,000 permutations within inhibition type (seed 42; plus-one correction). |
| ALM error analysis | 10 rank-10 fits to balanced correct, uninhibited training trials | Measured activity projected with Q, then movement regression; five session-grouped decoder folds and shuffled controls. Regional AUC: 26 bins in [−0.5, 0] s. |

## Acknowledgements

This work was supported by the Medical Research Council awards MC UU 00003/6 (to A.S.) and UKRI/MR/B000936/1. M.K. was supported by an Engineering and Physical Sciences Research Council (EPSRC) studentship within the University of Oxford and the IBM Computational Discovery DPhil programme. R.P. was supported by the Deutsche Forschungsgemeinschaft (DFG, German Research Foundation; Project-ID 424778381, TRR 295).

## Data availability

The source recordings are publicly available as Perich–Miller Dandiset 000688, version 0.250122.1735 [37], and Mesoscale Activity Map Dandiset 000363, version 0.231012.2129 [38]. The RNN analysis notebook retrieves the pretrained checkpoints and trajectories accompanying MARBLE [16] from Harvard Data-verse (https://doi.org/10.7910/DVN/KTE4PC). Processing scripts and session-inclusion lists are included with the analysis code.

## Code availability

Neuralign source code will be made publicly available at https://github.com/MiklosKralik/neuralign. Manuscript analyses and figure-generation scripts will be released at the same time at https://doi.org/10.5281/zenodo.24178673.

## Declarations

### Author contributions

Conceptualisation: *M.K, S.T, J.K, A.S, R.P*. Methodology: *M.K, J.K, R.P*. Software and analysis: *M.K, R.P*. Supervision: *J.K, R.P.C, A.S, R.P*. Funding acquisition: *M.K, A.S*. Writing—original draft: *M.K, R.P*. Writing—review and editing: *M.K, S.T, J.K, R.P.C, A.S, R.P*.

### Competing interests

The authors declare no competing interests.

### Ethics

This study reanalysed previously published datasets. No new animal experiments were performed for this work.

## Notes

### Competing Interest Statement

The authors have declared no competing interest.

